# Longitudinal pathway analysis using structural information with case studies in early type 1 diabetes

**DOI:** 10.1101/2022.06.21.497110

**Authors:** Maria K. Jaakkola, Anu Kukkonen-Macchi, Tomi Suomi, Laura L. Elo

## Abstract

We introduce a new method for Pathway Analysis of Longitudinal data (PAL), which is suitable for complex study designs, such as longitudinal data. The main advantages of PAL are the use of pathway structures and the suitability of the approach for study settings beyond currently available tools. We demonstrate the performance of PAL with three longitudinal datasets related to the early development of type 1 diabetes, involving different study designs and only subtle biological signals. Transcriptomic and proteomic data are represented among the test data. An R package implementing PAL is publicly available at https://github.com/elolab/PAL.

**Motivation:** Pathway analysis is a frequent step in studies involving gene or protein expression data, but most of the available pathway methods are designed for simple case versus control studies of two sample groups without further complexity. The few available methods allowing the pathway analysis of more complex study designs cannot use pathway structures or handle the situation where the variable of interest is not defined for all samples. Such scenarios are common in longitudinal studies with so long follow up time that healthy controls are required to identify the effect of normal aging apart from the effect of disease development, which is not defined for controls. PAL is the first available pathway method to analyse such high-investment datasets.

## Introduction

Pathway analysis is a routine step when analysing gene or protein expression data and many tools have been developed for it. Most of them are very simple and provide either a list of pathways with different activity between case and control sample groups (e.g. Tarca *et al*., 2009, Gu and Wang, 2013, Zyla *et al*., 2019, Fang *et al*., 2012, Dong *et al*., 2016, Sun *et al*., 2017), or pathway scores for each sample as such (e.g. Tomfohr *et al*., 2005, Hänzelmann *et al*., 2013, Foroutan *et al*., 2018, Jaakkola *et al*., 2018, Liu *et al*., 2017, Drier *et al*., 2013). However, many research projects involving transcriptomic or proteomic data have a more complex study design making the available pathway methods unsuited for them. In addition, the majority (e.g. all of the above mentioned examples) of the available pathway methods are designed and validated with transcriptomic data only.

Here we introduce a novel method for Pathway Analysis of Longitudinal data (PAL), which is designed for complex study designs. PAL can neutralise the effect of given confounding variables and identify pathways with significant associations with a user-defined main outcome. Both the variables to be neutralised and the main outcome variable can be either numeric or categorical, which supports the versatility of the method. PAL is suited to analyse longitudinal data and it can handle non-aligned time points, which is important in many studies involving long follow-up time. While longitudinal data is the most important application type for PAL, it can also be applied into other types of study settings, including simple ones, such as multiple categorical sample groups. The first main advantage of PAL is that it can estimate the significance of pathways related to outcome variables defined only for case samples (e.g. time-to-event or disease stage) and still control for relevant confound variables present for all samples (e.g. age in studies with long follow-up time). As far as we know, no other pathway method enables this. Another major advantage of PAL is its ability to use pathway structures, which is important as methods utilising pathway structures have been shown to outperform simple gene set approaches in the context of straightforward case versus control comparisons (Nguyen *et al*. 2019, Lim *et al*. 2020, Jaakkola and Elo 2016). In this study, we also show that besides transcriptomic data, PAL can be successfully applied to proteomic data, which further expands its potential applications.

There are few methods that allow the pathway analysis of longitudinal data. They include Attractor analysis of Boolean network of Pathway (ABP) (Sun *et al*., 2019), Longitudinal Linear Combination Test (LLCT) (Khodayari Moez *et al*., 2019), Time-Course Gene Set Analysis (TcGSA) (Hejblum *et al*., 2015), and Gene Set Enrichment Analysis (GSEA) for time series (Subramanian *et al*. 2005). ABP allows sophisticated post-processing of sample-specific pathway scores, but it has not been implemented into an R package, which hinders the ease of its usage. LLCT is a diverse tool building on the principles of mixed effects modelling and it can also be used with small sample size. TcGSA uses mixed effects modelling with maximum likelihood estimates to identify gene sets with significant variation over time. GSEA is a widely used pathway method that can take time points as phenotypes, though such continuous variables can not be combined with categorical ones. Despite the flexibility, usability and other benefits of these methods, none of them can be used to analyse data with a variable unavailable for control samples (e.g. disease stage or time-to-event), which makes them unsuited for such study settings. They also consider pathways as simple gene sets without further structural complexity.

Type 1 diabetes (T1D) is a complex autoimmune disease usually developing during childhood. As its progression is largely unknown, long follow up studies involving children with the genetic risk factors are needed. Indeed, many such longitudinal studies have been carried out or are ongoing, such as Type 1 Diabetes Prediction and Prevention (DIPP) (Kallionpää *et al*. 2014), Pathogenesis of Type 1 Diabetes - Testing the Hygiene Hypothesis (Diabimmune) (Kallionpää *et al*. 2019), and The Environmental Determinants of Diabetes in the Young (TEDDY) (Elding Larsson *et al*. 2014). They are considerable investments of time, money, and other resources, and suitable computational tools are required to fully utilise the large-scale omics data generated in them. However, such data are not straightforward to analyse as the normal age-related development of the children causes changes that can mask the delicate disease progression related biological signals. Appearance of certain T1D-related autoantibodies, called *seroconversion*, is the most reliable predictor of T1D (Chantal *et al*. 2018), but very little is known about processes leading to it. In this study, we used three publicly available longitudinal proteomic and transcriptomic datasets of children before and after seroconversion and healthy controls to demonstrate the utility of PAL for interpretation of these challenging real-life datasets.

## Results

Here we introduce a novel pathway method PAL, which allows the analysis of complex study designs. In the PAL workflow, the effects of user-defined confounding variables are neutralised at gene or protein level, pathway scores are calculated for all samples using pathway structures, and the significance of their association with a given main variable of interest is calculated using, by default, linear mixed effects models (Figure 1). A more detailed description of PAL is available in section Materials and methods. To demonstrate the utility of PAL for detecting significant pathways in respect to different variables of interest, we analysed three different datasets containing longitudinal data from prediabetic and healthy control children. In all these case studies, the normal age effect was first neutralised according to the healthy control samples using age as fixed effect and donor as random effect in the model fitting, and the significance for each pathway regarding 1) the time from seroconversion, and 2) the sample group (case vs control) was calculated using the variable of interest as fixed effect and donor as random effect in the model fitting.

**Figure 1:**
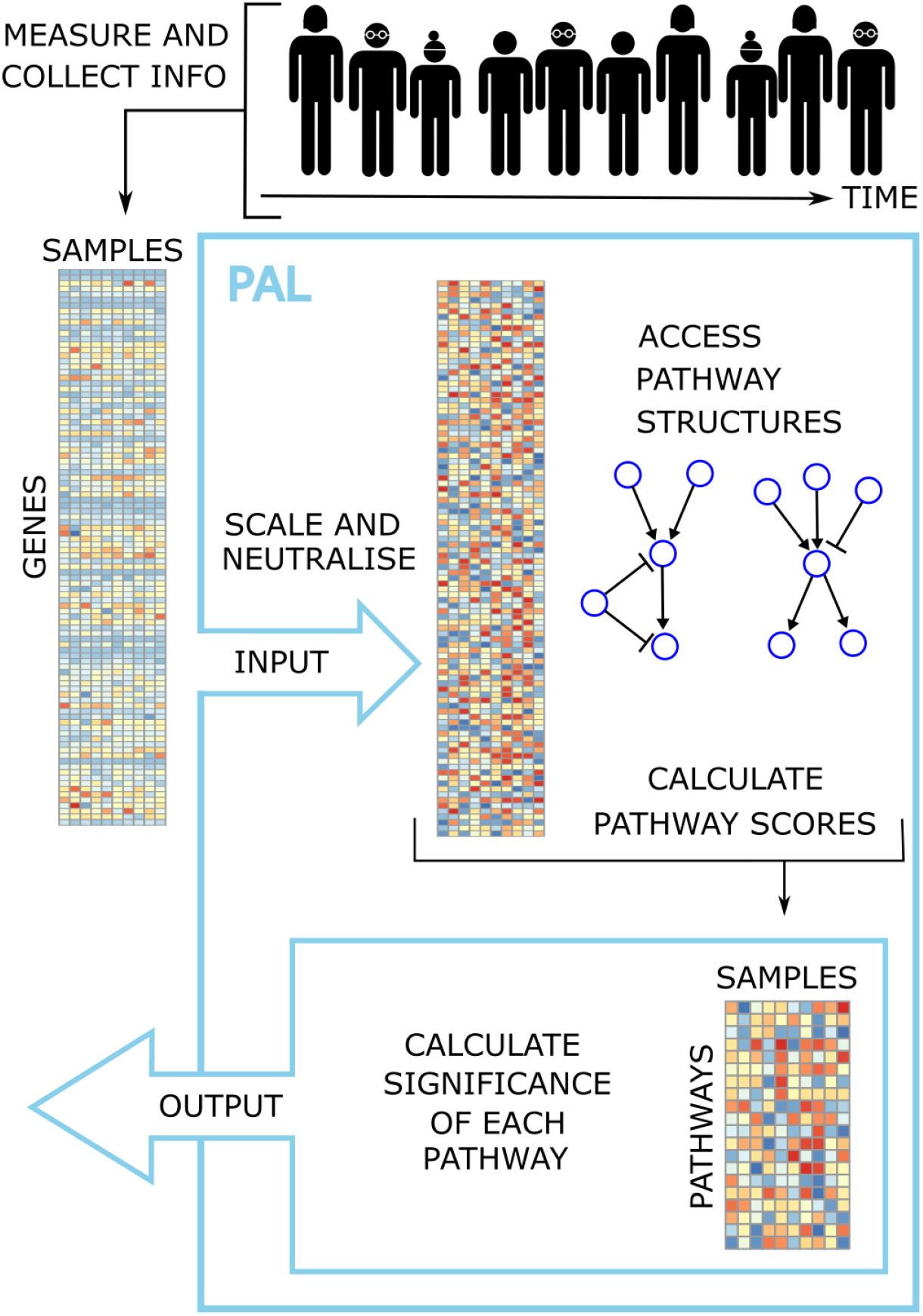
Schematic overview of the PAL workflow.

### Case study 1: Plasma proteomic data with long follow-up time

Application of PAL to the DAISY plasma proteomics data from 11 prediabetic and 10 healthy donors revealed nine pathways significantly associated with time from seroconversion after neutralising the age effect based on control samples (FDR < 0.05, Supplementary Table 1). Among these findings, pathway ‘Phenylalanine, tyrosine and tryptophan biosynthesis’ (KEGG id hsa00400) had a particularly clear time from seroconversion effect (Figure 2A).

**Figure 2.**
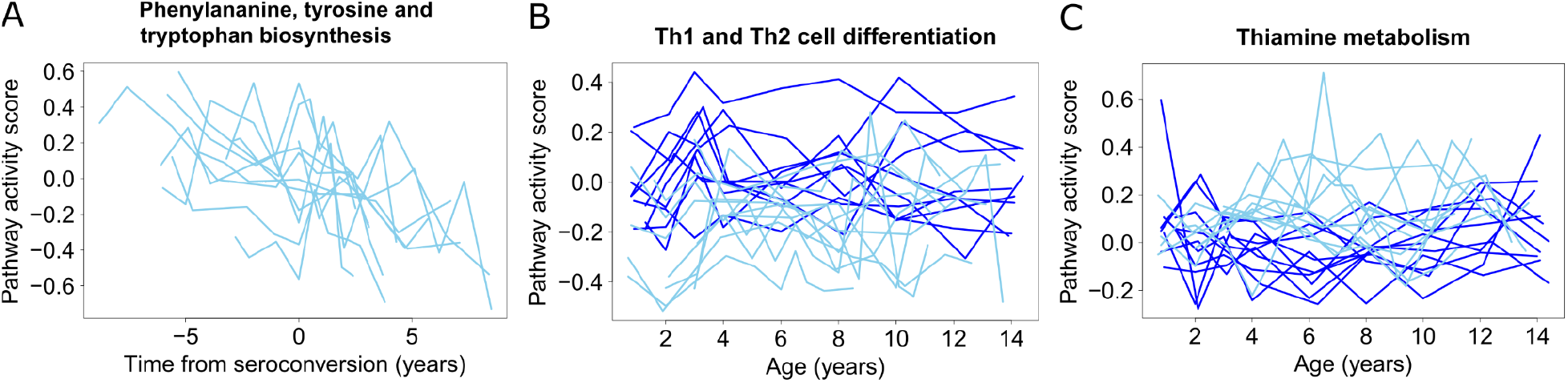
PAL results for pathways **(A)** Phenylalanine, tyrosine and tryptophan biosynthesis, **(B)** Th1 and Th2 cell differentiation, and **(C)** Thiamine metabolism. Pathway activity score estimated by PAL (y-axis) is illustrated as a function of time from seroconversion (A) or age (B and C) for only the case samples (A) or for the case and control samples (B and C). Each line represents one donor, with cases in light blue and controls in dark blue.

From the comparison between the prediabetic and healthy donors, 88 significant pathways were identified with PAL (FDR < 0.05, Supplementary Table 1). Of these pathways, 75 had higher activity in the healthy control subjects (e.g. ‘Th1 and Th2 cell differentiation’, Figure 2B) and 13 had higher activity in the prediabetic subjects (e.g. ‘Thiamine metabolism’, Figure 2C). Out of the 88 significant detections, 25 had sample group coefficient with absolute value > 0.1. These 25 strong detections included eight disease pathways, seven pathways related to metabolism of different biomolecules, five signalling pathways, and five other pathways, namely ‘Ferroptosis’, ‘Protein export’, ‘Th1 and Th2 cell differentiation’, ‘Nucleotide excision repair’, and ‘Mineral absorption’.

As seroconversion in T1D is still not fully understood and the underlying reasons leading to it are heterogeneous and likely include both genetic and environmental factors (Blanter *et al*. 2019), it is difficult to determine which pathways should be detected and which ones not. Several of the nine significant pathways associated with the time from seroconversion were related to metabolism of different biomolecules, which have been identified also in other studies. The strongest detection was ‘Phenylalanine, tyrosine and tryptophan biosynthesis’ and particularly changes related to phenylalanine and tyrosine have been found in the context of early stage T1D (Kostic *et al*. 2015, Sen *et al*. 2020), supporting also the significant pathways ‘Tyrosine metabolism’ and ‘Phenylalanine metabolism’. Similarly, the pathway ‘Bile secretion’ is supported by the literature (Lamichhane *et al*. 2021), while ‘Pancreatic secretion’ has obvious relevance. Peroxisome proliferator-activated receptors (PPARs) regulate processes like T cell differentiation and insulin secretion, supporting the significant detection of ‘PPAR signaling pathway’.

The strongest finding from the sample group comparison was ‘Alpha-Linoleic acid metabolism’, which is supported by Overgaard et al. as they showed the deregulation of diglyceride in early states of the type 1 diabetes progression (Overgaard *et al*. 2016). The second strongest detection was ‘Ferroptosis’, referring to iron-dependent regulated cell death, which was identified only 10 years ago and is currently under intensive study; it has already been linked to the development of metabolic diseases (Duan *et al*. 2021). The pathway ‘Th1 and Th2 cell differentiation’ has been strongly linked to development of T1D (Walker *et al*. 2016, Bradley *et al*. 1999). Besides ‘Alpha-Linoleic acid metabolism’, among the strong biomolecule metabolism pathway detections ‘Alanine, aspartate and glutamate metabolism’ and ‘Thiamine metabolism’ have also been linked to T1D (Sen *et al*. 2020, Pácal *et al*. 2014). In addition, different glycans and ascorbate are related to immune system, supporting the detected pathways ‘N-Glycan biosynthesis’, ‘Glycosaminoglycan biosynthesis - chondroitin sulfate / dermatan sulfate’, and ‘Ascorbate and aldarate metabolism’. Several of the strong signalling pathway detections are also related to immune system, as ‘RIG-I-like receptor signaling pathway’ works in defence against virus infections, ‘Rap1 signaling pathway’ has been shown to regulate T cell response (Katagiri *et al*. 2002), and ‘IL-17 signaling pathway’ recruits immune cells to the site of inflammation and it has been linked to different autoimmune diseases (Hu *et al*. 2011). ‘Notch signaling pathway’ has very diverse functions including regulating apoptosis and pancreatic development and its potential role in development of T1D has been discussed (Kim *et al*. 2010). The eight disease related strong pathway findings from the sample group comparison included autoimmune diseases (‘Autoimmune thyroid disease’ and ‘Type I diabetes mellitus’), rejection reaction related diseases (‘Graft-versus-host disease’, ‘Allograft rejection’, and ‘Endocrine resistance’), and infection pathways (‘Salmonella infection’ and ‘African trypanosomiasis’). While there is unlikely to be direct causality between the early stage T1D and these diseases (excluding T1D itself), they share some mechanisms or characteristics, which can explain the detection of these pathways.

Finally, we wanted to compare the PAL findings to the findings in the original study. Notably, as no other pathway tools can directly analyse this study design, the only alternative to PAL was to do the analysis at the protein level and then use the detected proteins as an input for a standard pathway analysis method. In the original publication (Liu *et al*., 2018), only two significantly differentially expressed proteins were detected at FDR of 0.05 between the prediabetic and healthy subjects. Therefore, we used a more relaxed FDR cutoff of 0.1, with which 29 differentially expressed proteins were identified. Database for Annotation, Visualization and Integrated Discovery (DAVID) analysis of them did not identify any significant KEGG pathways. This highlights the utility of PAL for discovering significant pathways from challenging datasets with complex designs and only delicate signals.

### Case study 2: Transcriptomic time series data from young children

The Diabimmune data (Kallionpää *et al*. 2019) containing gene expression measurements from seven prediabetic and eight healthy children were analysed using PAL separately for the peripheral blood mononuclear cells (PBMCs), CD4+ T cells, and CD8+ T cells. Only one pathway in PBMC data, namely ‘Lysosome’, was significantly associated with time from seroconversion after neutralising the age effect based on control samples (FDR < 0.05, Supplementary Table 1).

In the comparison between the prediabetic and healthy control subjects, 28 pathways were significant in the PBMC samples, 17 in the CD4+ T cells, and 99 in the CD8+ T cells (Supplementary Table 1). This is in line with the original study (Kallionpää *et al*. 2019), where early changes in prediabetes already before seroconversion were reported particularly in the CD8+ T cells. Interestingly, the significant pathways from CD4+ T cells and CD8+ T cells were almost exclusively downregulated in case samples, whereas in the PBMC data 16 of the significant pathways were downregulated and 12 were upregulated in the case samples. Among the significant pathways, four had absolute sample group coefficient > 0.1 in PBMC data, three in CD4+ T cell data, and 25 in CD8+ T cell data (Supplementary Table 1). Pathways ‘Spliceosome’, ‘Mitophagy - animal’, and ‘Circadian rhythm’ were among these strongest detections in both PBMC and CD8+ T cells, and they were all downregulated in case samples in both cell types (Figure 3).

**Figure 3.**
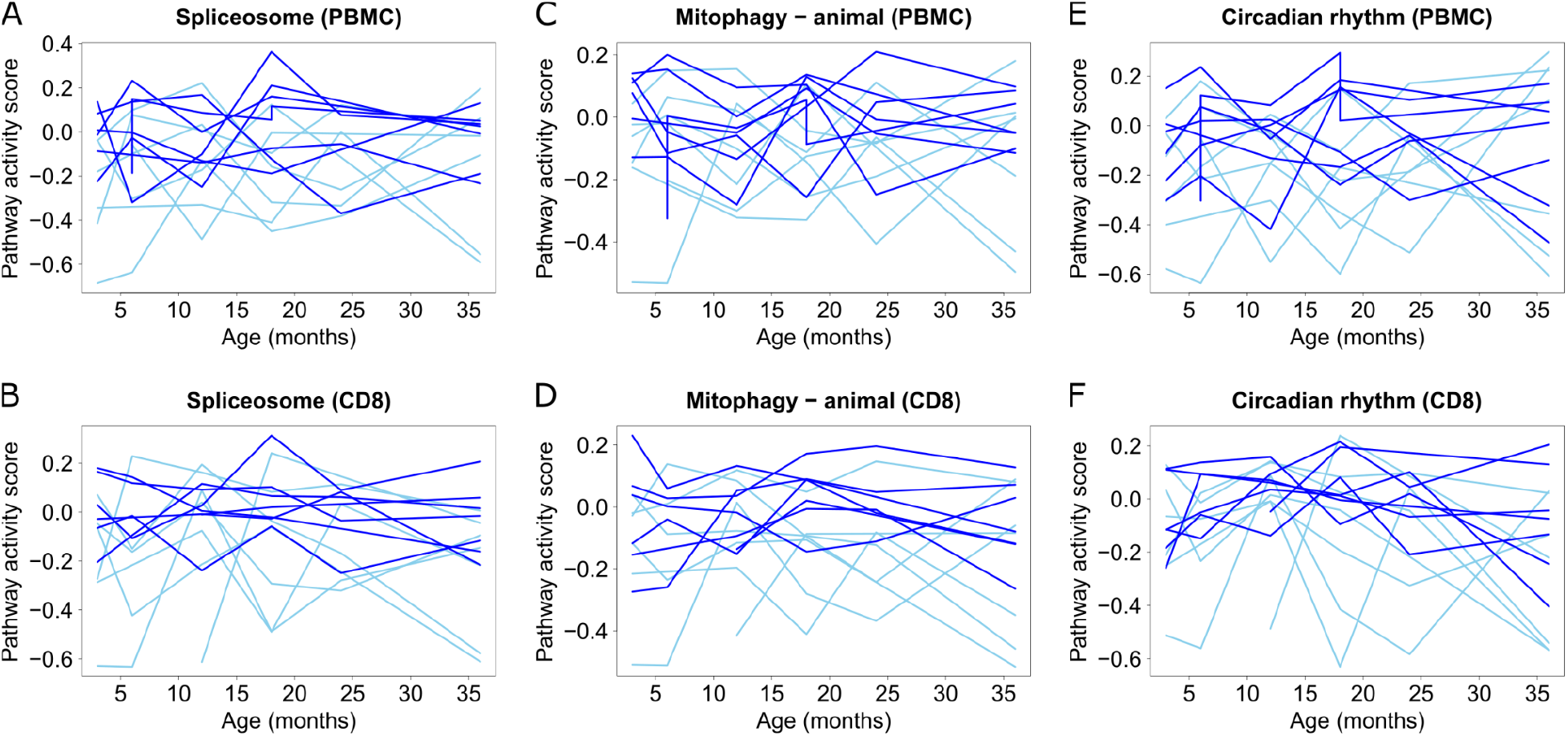
Pathway activity score (y-axis) as function of age (x-axis) of **(A-B)** Spliceosome, **(C-D)** Mitophagy - animal, and **(E-F)** Circadian rhythm in PBMC (upper panel) and in CD8+ T cells (lower panel). Case individuals are indicated with light blue lines and control individuals with dark blue lines.

The 25 strongest detections with the absolute sample group coefficient > 0.1 from CD8+ T cells (Supplementary Table 1) included several pathways that have been linked to the destruction of beta cells (‘Autophagy - other’, ‘Proteasome’, ‘Mitophagy - animal’, ‘JAK-STAT signaling pathway’, ‘Protein processing in endoplasmic reticulum’) (Lambelet *et al*. 2018, Thomaidou *et al*. 2018, Sidarala *et al*. 2020, Gurzov *et al*. 2016, Vig *et al*. 2021), but it is unclear if such processes are altered already at this early stage of the disease development. Three of the strong findings (‘Autophagy - other’, ‘Proteasome’, and ‘Mitophagy - animal’) are related to within-cell homeostasis and one (‘Apoptosis - multiple species’) to tissue/organ-level homeostasis. The ‘JAK-STAT signaling pathway’ is also related to apoptosis and the ‘B cell receptor signaling pathway’ to autoimmunity (Rawlings *et al*. 2017). Changes in citrate acid cycle have been identified in children who will develop T1D in the future (Oresic *et al*. 2008), which supports two of the strong detections (‘2-Oxocarboxylic acid metabolism’ and ‘Citrate cycle (TCA cycle)’), as 2-Oxocarboxylic acid is part of the TCA cycle. Besides the TCA cycle, ‘Oxidative phosphorylation’ is related to the ATP production. The most interesting observation among the strongest disease specific pathways was the presence of three pathways related to cardiomyopathy (‘Arrhythmogenic right ventricular cardiomyopathy’, ‘Viral myocarditis’, and ‘Dilated cardiomyopathy’). While increased risk of cardiomyopathy among T1D patients is strongly supported in the literature (Ritchie *et al*. 2017), this connection with early stage disease was surprising.

Eight of the significant detections from the PBMC data were also identified from the DAISY proteomics data. Among these eight pathways, four (‘Protein processing in endoplasmic reticulum’, ‘Autophagy - animal’, ‘Non-small cell lung cancer’, and ‘Epstein-Barr virus infection’) were downregulated in case samples of both dataset and one (‘Biosynthesis of cofactors’) was upregulated in them.

To compare the PAL findings to the findings in the original study (Kallionpää *et al*. 2019), we considered the genes reported to be differentially expressed between the sample groups across all timepoints (FDR < 0.05) and used the DAVID pathway tool. With this, we did not identify any significant KEGG pathways at FDR of 0.05 from the CD8+ T cells or the CD4+ T cells, whereas 17 pathways were detected from the PBMCs (Supplementary Table 1). This is an interesting observation as PAL detected the largest number of pathways from the CD8+ T cells. One DAVID detection from the PBMC samples (‘Epstein-Barr virus infection’) overlapped with the corresponding PAL detections, and four (‘Th1 and Th2 cell differentiation’, ‘Th17 cell differentiation’, ‘Epstein-Barr virus infection’, and ‘Type I diabetes mellitus’) overlapped with the PAL detections from the CD8+ T cells.

### Case study 3: Transcriptomic longitudinal data with long follow up time

Finally, we applied PAL to the BabyDiet data (Ferreira *et al*. 2014) containing longitudinal transcriptomic PBMC data measured from 22 prediabetic and 87 healthy control children. PAL detected seven significant pathways associated with the time from seroconversion after neutralising the age effect based on the control samples (FDR < 0.05, Supplementary Table 1), but the absolute coefficients were small indicating that the effect was relatively weak despite being significant. All of the seven significant pathways had increasing trend over time from seroconversion and four out of the seven (‘Thiamine metabolism’, ‘Lysine degradation’, ‘Purine metabolism’, and ‘Primary bile acid biosynthesis’) were related to metabolism of different biomolecules. Relatedly, the detected pathway ‘One carbon pool by folate’ supports biosynthesis and homeostasis of several amino acids (Ducker and Rabinowitz 2017).

PAL identified seven significant pathways with altered activity between the sample groups, but again, the absolute coefficients were relatively small (Supplementary Table 1). All of the seven significant pathways were upregulated in the case samples. The strongest detection was the pathway ‘Phagosome’ while all the other significant findings were related to metabolism of different biomolecules.

Time from seroconversion had a significant effect on two pathways related to the genome and on five pathways associated with metabolism of different biomolecules. The pathway ‘One carbon pool by folate’ was the strongest among the latter. One-carbon metabolism mediated by the folate cofactor supports methionine homeostasis (Ducker and Rabinowitz 2017), and methionine has been associated with early stage T1D (Overgaard *et al*. 2016). In addition, Pácal *et al*. have studied connections between altered thiamine metabolism and diabetes (Pácal *et al*. 2014), Li *et al*. have provided a clear overview of roles of different amino acids, including lysine, in immune system (Li *et al*. 2007), and Ahmad and Haeusler have evaluated the roles of bile acids in glucose metabolism and insulin signalling (Ahmad and Haeusler 2019), supporting the detected pathways ‘Thiamine metabolism’, ‘Lysine degradation’, and ‘Primary bile acid biosynthesis’, respectively.

From the sample group comparison, six of the seven significant detections were related to metabolism of different biomolecules. The only exception was pathway ‘Phagosome’, which is part of the immune system. Among the detected biomolecule metabolism pathways, ‘Alanine, aspartate and glutamate metabolism’ has also been identified in another study of early T1D (Sen *et al*. 2020), while ‘Retinol metabolism’ (retinol is a type of vitamin A) plays a role in immune system (Kim *et al*. 2015) and two of the detections (‘Galactose metabolism’ and ‘Pentose and glucuronate interconversions’) are related to sugar metabolism. The detected ‘Biosynthesis of cofactors’ is a large pathway inducing the biosynthesis of multiple cofactors, including retinol. Interestingly, pathways ‘Alanine, aspartate and glutamate metabolism’ and ‘Biosynthesis of cofactors’ were also detected from both the DAISY and the Diabimmune PBMC datasets when the sample groups were compared. In addition, ‘Phagosome’ was detected from the DAISY and ‘Retinol metabolism’ from the Diabimmune PBMC data.

## Discussion and conclusions

Here we introduced a new method for pathway analysis of longitudinal data PAL and demonstrated its performance with challenging longitudinal data with only moderate biological signals. Even from these datasets, PAL identified several relevant pathways that remained undetected in the original studies but were well supported by the literature. Particularly pathways related to metabolism of different biomolecules were well represented among the detections.

The tested datasets were of different origin (proteomics, RNA-sequencing, and gene expression microarrays of serum, PBMCs, CD4+ T cells, or CD8+ T cells). This, together with the heterogeneity of T1D and the very early disease stage, likely explains the modest overlaps between the detected pathways from the different datasets. When detecting pathways significantly associated with time from seroconversion, PAL detected the largest number of pathways in the DAISY dataset, and those detections included many processes associated with the development of T1D in the literature, suggesting the utility of proteomics for studying the early stage development of T1D. In general, PAL detected more significant pathways from the sample group comparison than from the time from seroconversion analysis.

The benefits of breaking the model fitting into two parts (neutralisation step at gene/protein level and significance step at pathway level) are: 1) the main variable of interest (e.g. disease stage) may not be defined for control samples, but with this approach they can still be utilised, 2) while the significance step has to be done at pathway level, doing the neutralisation already at gene/protein level supports the robustness of the approach, and 3) using only the control samples for neutralisation does not mask the possibly altered effect of these variables in the case samples. In addition, PAL provides a very flexible user interface and utilises pathway structures instead of simple gene sets.

The main limitation of PAL is its inability to neutralise categorical features that are only partially shared between the sample groups. A toy example of a partially shared feature is ethnicity when cases and controls include ethnicities ‘caucasian’ and ‘hispanic’, but only cases have also some ‘asian’ samples. If both sample groups would include all three groups, there would be no problem. However, such an imbalanced study setting can also be seen as a weakness of the original study itself. This study suffers from difficulty of validation. As T1D is widely studied, but heterogeneous and not that well understood, very many detected pathways can be linked to its development, but there is no gold standard available. However, datasets like these are the ones that most benefit from PAL analysis, which makes them realistic case studies for demonstration purposes.

To summarise, PAL is a novel pathway method suitable for analysing datasets with complex study design, such as longitudinal data. It is simple to use, can be applied on transcriptomic or proteomic data, uses pathway structures, detects relevant pathways even from challenging data, and allows the analysis of study designs that cannot be analysed with the other currently available pathway methods.

## Materials and methods

### The PAL algorithm

PAL requires normalised expression data and sample group information (dummy values can be used if the study design has no sample groups) as mandatory inputs, but further information about the samples enables analysis of complex study designs. Based on the additional information, the effect of some variables (e.g. normal aging) can be neutralised already at the gene/protein level. Then PAL will automatically access KEGG pathways via API and determines sample-specific pathway scores utilising pathway structures with the Pathway Analysis for Sample-level Information (PASI) method (Jaakkola *et al*., 2018). Finally, if sufficient input is provided, PAL calculates the significance of a given variable (e.g. disease stage) to the pathway activity/deregulation scores while considering the effect of given variables, such as donor, in the model fitting.

The effect of a given confounding variable is neutralised separately for each gene/protein prior to the pathway analysis. The default neutralisation step starts with fitting a linear mixed effects model to the control samples using function lmer from the R package lme4 (Bates *et al*., 2015). By default, the given confounding variables to be neutralised are used as fixed effects and given variables (e.g. donor) as random effects. Then the estimated coefficients multiplied by the corresponding observed values are subtracted from the expression values to neutralise their effect. The variables to be neutralised can be either numeric (e.g. age or bmi), or categorical (e.g. smoking status) and many of them can be provided simultaneously. The end-user can also provide a custom formula used in the model fitting, if the default model is unsuited for their study design, and the linear mixed effects model itself can be replaced with either robust linear model (function rlm from R package MASS (Venables and Ripley, 2002)) or robust linear mixed effects model (function rlmm from R package robustlmm (Koller 2016)) using the relevant arguments. However, the robust mixed effect model is slow to fit, which limits its usage (see PAL R package documentation in GitHub for further information). Notably, if the number of variables in the model is high and the sample size is not, the risk of overfitting is considerable. With PASI, it is recommended to use at least 10-15 control samples (Jaakkola *et al*., 2018), but multiple variables to be neutralised likely require a higher number in PAL analysis. The pathway scores are calculated from these processed gene/protein expression values using PASI. The output is a matrix of pathway scores with samples as columns and pathways as rows. The pathway scores can represent either activity (default) or deregulation of the analysed pathways.

Finally, the significance of each pathway is calculated from the pathway scores with respect to the given outcome of interest. Notably, not all samples necessarily have the variable of interest (e.g. seroconversion time is missing from control samples). By default, a linear mixed effects model with a given variable of interest as fixed effect and given random effects (e.g. donor) is fitted for each pathway using the function lmer from the R package lme4. However, using the corresponding arguments (see the R package documentation for PAL in GitHub), the user can choose to use either robust linear model with only fixed effect (rlm) or robust linear mixed effect model (rlmm) instead of lmer, and/or give different model formula to be fitted. In each pathway, the p-value for the given main variable of interest is calculated by first fitting the model and extracting the coefficient describing the effect of the main variable, and then randomly sampling the main variable over the samples multiple times (n=1000 in this study). Then it is calculated how often the original coefficient has value greater than the corresponding value from a model with a randomly sampled variable and those frequencies are converted into p-value estimates. The resulting p-values are then further converted into false discovery rates (FDR) using function p.adjust with the method argument “fdr”, and finally the p-values, the FDR values, and the coefficients are returned for all pathways together with the sample specific pathway scores. It is possible to provide variables to be neutralised, outcome variable of interest, both, or neither. The PAL R package documentation in GitHub contains further instructions about the usage.

### Datasets

We demonstrate the accuracy and utility of PAL with three publicly available datasets involving longitudinal proteomics or transcriptomics data.

DAISY dataset was downloaded from ProteomeXchange (Deutsch *et al*., 2020) with accession number PXD007884. It contains longitudinal plasma proteomics data across nine time points from 11 prediabetic and 10 healthy 0-15 years old donors, measured using TMT-based quantitative mass spectrometry (Liu *et al*., 2018). The dataset was preprocessed similarly as in (Välikangas *et al*. 2021). The protein identifiers were converted to corresponding Entrez gene identifiers using R package biomaRt (version 2.48.3) (Durinck *et al*. 2009).

Diabimmune dataset was downloaded from European Genome-phenome Archive with accession number EGAC00001001443. It contains longitudinal transcriptomics data across five time points (3, 6, 12, 24, and 36 months of age) from 7 seroconverted and 8 healthy control children (Kallionpää *et al*., 2019). All the sample donors have increased genetic risk of T1D.

Peripheral blood mononuclear cells (PBMC), CD4+ T cells, and CD8+ T cells were measured separately using RNA-sequencing. The data was normalised using the Trimmed Mean of M-values (TMM) method.

BabyDiet dataset was downloaded from Array Express with accession number MTAB1724. It contains longitudinal transcriptomics data of PBMC samples across 1 to 12 time points (median 4) from 22 seroconverted and 87 healthy control children, measured using the the Affymetrix GeneChip Human Gene 1.1 ST microarray (Ferreira *et al*. 2014). The readily available preprocessed data was used. While the age range of the children was wide, ranging from young babies to 10 years of age, the majority of the samples were from children under 3 years and we removed outlier samples according to age prior to the analyses. The cutoff for an outlier was defined using interquartile range (IQR) criteria, in which normal range was defined from 1st quartile - 1.5*IQR to 3rd quartile + 1.5*IQR, where IQR = (3rd quartile - 1st quartile). One case donor and 15 control donors had only one sample and these samples without longitudinal aspect were also removed prior to the analysis.

### Statistical analysis

The baseline pathway analysis was done by using the Database for Annotation, Visualization and Integrated Discovery (DAVID) tool (2021 update) on differentially expressed genes/proteins identified in the original studies introducing the data. Notably, no such differentially expressed genes were available from the BabyDiet study, so the baseline analysis is missing for that dataset. All analysed genes/proteins were used as a background for DAVID. From the DAVID results, statistically significant KEGG pathways were considered. In all analyses, a finding was considered as statistically significant if it had FDR below 0.05, unless otherwise stated.

## Author contributions

MKJ participated in designing, implementing, and documenting the method, planning the study design, running the analyses, interpreting the results, and preparing the manuscript. AK-M participated in interpreting the results and preparing the manuscript. TS participated in testing and documenting the R package, supervising the project, and preparing the manuscript. LLE conceived the study, participated in the study design and preparing the manuscript, and supervised the project. All the authors approved the final manuscript.

## References

Ahmad, T.R. and Haeusler, R.A. (2019) Bile acids in glucose metabolism and insulin signalling—mechanisms and research needs. Nature Reviews Endocrinology, 15, 701–712.

Bates D., M{\”a}chler M., Bolker B., and Walker S. (2015) Fitting Linear Mixed-Effects Models Using lme4. Journal of Statistical Software, 67, 1–48.

Blanter, M., Sork, H., Tuomela, S., and Flodström-Tullberg, M. (2019) Genetic and environmental interaction in type 1 diabetes: a relationship between genetic risk alleles and molecular traits of enterovirus infection? Current Diabetes Reports, 19, 1–14.

Bradley, L.M., Asensio, V.C., Schioetz, L.K., Harbertson, J., Krahl, T., Patstone, G., Woolf, N., Campbell, I.L., and Sarvetnick, N. (1999) Islet-specific Th1, but not Th2, cells secrete multiple chemokines and promote rapid induction of autoimmune diabetes. The Journal of Immunology, 162, 2511–2520.

Chantal, M., Lahesmaa, R., Bonifacio, E., Achenbach, P., and Tree, T. (2018) Immunological biomarkers for the development and progression of type 1 diabetes. Diabetologia, 61, 2252–2258.

Deutsch, E.W., Bandeira, N., Sharma, V., Perez-Riverol, Y., Carver, J.J., Kundu, D.J., García-Seisdedos, D., Jarnuczak, A.F., Hewapathirana, S., Pullman, B.S., and Wertz, J. (2020) The ProteomeXchange consortium in 2020: enabling ‘big data’ approaches in proteomics, Nucleic Acids Research, 48, D1145–D1152.

Dong, X., Hao, Y., Wang, X., and Tian, W. (2016) LEGO: a novel method for gene set over-representation analysis by incorporating network-based gene weights. Scientific reports, 6, 1–17.

Drier, Y., Sheffer, M., and Domany, E. (2013) Pathway-based personalized analysis of cancer. Proceedings of the National Academy of Sciences, 110, 6388–6393.

Duan, J.-Y., Lin X.., Xu F., Shan S.-K., Guo B., Li F.-X.-Z.., Wang Y., Zheng M.-H., Xu Q.-S., Lei L.-M. et al. (2021) Ferroptosis and its potential role in metabolic diseases: a curse or revitalization? Frontiers in Cell and Developmental Biology, 9.

Ducker, G.S. and Rabinowitz, J.D. (2017) One-carbon metabolism in health and disease. Cell metabolism, 25, 27–42.

Durinck, S., Spellman, P.T., Birney, E., and Huber, W. (2009) Mapping identifiers for the integration of genomic datasets with the R/Bioconductor package biomaRt. Nature protocols, 4, 1184–1191.

Elding Larsson, H.E., Vehik, K., Gesualdo, P., Akolkar, B., Hagopian, W., Krischer, J., Lernmark, Å., Rewers, M., Simell, O., She, J.X., and Ziegler, A. (2014) Children followed in the TEDDY study are diagnosed with type 1 diabetes at an early stage of disease. Pediatric diabetes, 15, 118–126.

Fang, Z., Tian, W., and Ji, H. (2012) A network-based gene-weighting approach for pathway analysis. Cell research, 22, 565–580.

Ferreira, R.C., Guo, H., Coulson, R.M., Smyth, D.J., Pekalski, M.L., Burren, O.S., Cutler, A.J., Doecke, J.D., Flint, S., McKinney, E.F., and Lyons, P.A. (2014) A type I interferon transcriptional signature precedes autoimmunity in children genetically at risk for type 1 diabetes. Diabetes, 63, 2538--2550.

Foroutan, M.B., Dharmesh D., Lyu R., Horan, K., Cursons, J., and Davis, M.J. (2018) Single sample scoring of molecular phenotypes. BMC bioinformatics, 19, 1–10.

Gu, Z. and Wang, J. (2013) CePa: an R package for finding significant pathways weighted by multiple network centralities. Bioinformatics, 29, 658–660.

Gurzov, E.N., Stanley, W.J., Pappas, E.G.,Thomas, H.E., and Gough, D.J. (2016) The JAK/STAT pathway in obesity and diabetes. The FEBS journal, 283, 3002–3015.

Hänzelmann, S., Castelo, R., and Guinney, J. (2013) GSVA: gene set variation analysis for microarray and RNA-seq data. BMC bioinformatics, 14, 1–15.

Hejblum, B.P., Skinner, J., and Thiébaut, R. (2015) Time-Course Gene Set Analysis for Longitudinal Gene Expression Data. PLOS Computational Biology, 11, e1004310.

Hu, Y., Shen, F., Crellin, N.K., and Ouyang, W. (2011) The IL-17 pathway as a major therapeutic target in autoimmune diseases. Annals of the New York Academy of Sciences, 1217, 60–76.

Jaakkola, M.K. and Elo, L.L. (2016) Empirical comparison of structure-based pathway methods. Briefings in bioinformatics, 17, 336–345.

Jaakkola, M.K., McGlinchey, A.J., Klen, R., and Elo, L.L. (2018) PASI: A novel pathway method to identify delicate group effects. PloS one, 13, e0199991.

Kallionpää, H., Elo, L.L., Laajala, E., Mykkänen, J., Ricaño-Ponce, I., Vaarma, M., Laajala, T.D., Hyöty, H., Ilonen, J., Veijola, R., and Simell, T. (2014) Innate immune activity is detected prior to seroconversion in children with HLA-conferred type 1 diabetes susceptibility. Diabetes, 63, 2402–2414.

Kallionpää, H., Somani, J., Tuomela, S., Ullah, U., De Albuquerque, R., Lönnberg, T., Komsi, E., Siljander, H., Honkanen, J., Härkönen, T., and Peet, A. (2019) Early detection of peripheral blood cell signature in children developing b-cell autoimmunity at a young age. Diabetes, 68, 2024–2034.

Katagiri, K., Hattori, M., Minato, N., and Kinashi, T. (2002) Rap1 functions as a key regulator of T-cell and antigen-presenting cell interactions and modulates T-cell responses. Molecular and cellular biology, 22, 1001–1015.

Khodayari Moez, E., Hajihosseini, M., Andrews, J.L., and Dinu, I. (2019) Longitudinal linear combination test for gene set analysis. BMC bioinformatics, 20, 1–19.

Kim, M.H., Taparowsky, E.J., and Kim, C.H. (2015) Retinoic acid differentially regulates the migration of innate lymphoid cell subsets to the gut. Immunity, 43, 107–119.

Kim, W., Shin, Y.-K., Kim, B.-J., and Egan, J.M. (2010) Notch signaling in pancreatic endocrine cell and diabetes. Biochemical and biophysical research communications, 392, 247–251.

Koller M. (2016) robustlmm: An R Package for Robust Estimation of Linear Mixed-Effects Models. Journal of Statistical Software, 75, 1–24.

Kostic, A.D., Gevers, D., Siljander, H., Vatanen, T., Hyötyläinen, T., Hämäläinen, A.-M., Peet, A., Tillmann, V., Pöhö, P., Mattila, I. et al. (2015) The dynamics of the human infant gut microbiome in development and in progression toward type 1 diabetes. Cell host & microbe, 17, 260–273.

Lambelet, M., Terra, L.F., Fukaya, M., Meyerovich, K., Labriola, L., Cardozo, A.K., and Allagnat, F. (2018) Dysfunctional autophagy following exposure to pro-inflammatory cytokines contributes to pancreatic β-cell apoptosis. Cell death & disease, 9, 1–15.

Lamichhane, S., Sen, P., Dickens, A.M., Alves, M.A., Karkonen, T., Honkanen, J., Vatanen, T., Xavier, R.J., Hyotylainen, T., Knip, M. et al. (2021) Dynamics of gut microbiome-mediated bile acid metabolism in progression to islet autoimmunity. medRxiv.

Li, P., Yin, Y.L., Li, D., Kim, S.W., and Wu, G. (2007) Amino acids and immune function. British Journal of Nutrition, 98, 237–252.

Lim, S., Lee, S., Jung, I., Rhee, S., and Kim, S. (2020) Comprehensive and critical evaluation of individualized pathway activity measurement tools on pan-cancer data. Briefings in bioinformatics, 21, 36–46.

Liu, C., Lehtonen, R., and Hautaniemi, S. (2017) PerPAS: topology-based single sample pathway analysis method. IEEE/ACM transactions on computational biology and bioinformatics, 15, 1022–1027.

Liu, C.W., Bramer, L., Webb-Robertson, B.J., Waugh, K., Rewers, M.J., and Zhang, Q. (2018) Temporal expression profiling of plasma proteins reveals oxidative stress in early stages of Type 1 Diabetes progression. Journal of Proteomics, 172, 100–110.

Nguyen, T.-M., Shafi, A., Nguyen, T., and Draghici, S. (2019) Identifying significantly impacted pathways: a comprehensive review and assessment. Genome biology, 20, 1–15.

Oresic, M., Simell, S., Sysi-Aho, M., Nanto-Salonen, K., Seppänen-Laakso, T., Parikka, V., Katajamaa, M., Hekkala, A., Mattila, I., Keskinen, P. et al. (2008) Dysregulation of lipid and amino acid metabolism precedes islet autoimmunity in children who later progress to type 1 diabetes. The Journal of experimental medicine, 205, 2975–2984.

Overgaard, A.J., Kaur, S. and Pociot, F. (2016) Metabolomic biomarkers in the progression to type 1 diabetes. Current diabetes reports, 16, 1–5.

Pácal, L., Kuricová, K., and Kaňková, K. (2014) Evidence for altered thiamine metabolism in diabetes: Is there a potential to oppose gluco-and lipotoxicity by rational supplementation? World journal of diabetes, 5, 288.

Rawlings, D.J., Metzler, G., Wray-Dutra, M., and Jackson, S.W. (2017) Altered B cell signalling in autoimmunity. Nature reviews Immunology, 17, 421–436.

Ritchie, R.H., Zerenturk, E.J., Prakoso, D., and Calkin, A.C. (2017) Lipid metabolism and its implications for type 1 diabetes-associated cardiomyopathy. Journal of molecular endocrinology, 58, R225–R240.

Sen, P., Dickens, A.M., López-Bascón, M.A., Lindeman, T., Kemppainen, E., Lamichhane, S., Rönkkö, T., Ilonen, J., Toppari, J., Veijola, R., and Hyöty, H. (2020) Metabolic alterations in immune cells associate with progression to type 1 diabetes. Diabetologia, 63, 1017–1031.

Sidarala, V., Pearson, G.L., Parekh, V.S., Thompson, B., Christen, L., Gingerich, M.A., Zhu, J., Stromer, T., Ren, J., Reck, E.C. et al. (2020) Mitophagy protects β cells from inflammatory damage in diabetes. JCI insight, 5.

Subramanian, A., Tamayo, P., Mootha, V.K., Mukherjee, S., Ebert, B.L., Gillette, M.A., Paulovich, A., Pomeroy, S.L., Golub, T.R., Lander, E.S., Mesirov, J.P. et al. (2005) Gene set enrichment analysis: a knowledge-based approach for interpreting genome-wide expression profiles. Proceedings of the National Academy of Sciences, 102, 15545--15550.

Sun, D., Liu, Y., Zhang, X.S., and Wu, L.Y. (2017) NetGen: a novel network-based probabilistic generative model for gene set functional enrichment analysis. BMC systems biology, 11, 61–74.

Sun, S., Yu, X., Sun, F., Tang, Y., Zhao, J., and Zeng, T. (2019) Dynamically characterizing individual clinical change by the steady state of disease-associated pathway. BMC bioinformatics, 20, 1–12.

Tarca, A.L., Draghici, S., Khatri, P., Hassan, S.S., Mittal, P., Kim, J., Kim, C.J., Kusanovic, J.P., and Romero, R. (2009) A novel signaling pathway impact analysis. Bioinformatics, 25, 75–82.

Thomaidou, S., Zaldumbide, A., and Roep, B.O. (2018) Islet stress, degradation and autoimmunity. Diabetes, Obesity and Metabolism, 20, 88–94.

Tomfohr, J., Lu, J., and Kepler, T.B. (2005) Pathway level analysis of gene expression using singular value decomposition. BMC bioinformatics, 6, 1–11.

Välikangas, T., Suomi, T., Chandler, C.E., Scott, A.J., Tran, B.Q., Ernst, R.K., Goodlett, D.R., and Elo, L.L. (2021) Enhanced longitudinal differential expression detection in proteomics with robust reproducibility optimization regression. Preprint at bioRxiv

Venables, W.N. and Ripley, B.D. (2002) Modern Applied Statistics with S, Springer, New York.

Vig, S., Lambooij, J.M., Zaldumbide, A., and Guigas, B. (2021) Endoplasmic Reticulum-Mitochondria Crosstalk and Beta-Cell Destruction in Type 1 Diabetes. Frontiers in immunology, 12, 1271.

Walker, L.S.K. and von Herrath, M. (2016) CD4 T cell differentiation in type 1 diabetes. Clinical & Experimental Immunology, 183, 16–29.

Zyla, J., Marczyk, M., Domaszewska, T., Kaufmann, S.H.E., Polanska, J., and Weiner, J. 3rd (2019) Gene set enrichment for reproducible science: comparison of CERNO and eight other algorithms. Bioinformatics, 24, 5146–5154.

